# Microbiota-sensitive glial and metabolic programs define a critical window in early postnatal brainstem development

**DOI:** 10.64898/2026.07.13.738238

**Authors:** Cristina Cuesta-Marti, Caoimhe M.K. Lynch, Maya Davies, Indira Venkatesh, Austeja Baleviciute, Jessica Hincks, Lars Wilmes, Julia Eckenberger, Benjamin Valderrama, Jade A. Lawless, Gerard M. Moloney, Subrata Ghosh, Gerard Clarke, John F. Cryan, Lily Keane

**Affiliations:** APC Microbiota Ireland, University College Cork, Ireland; Department of Anatomy & Neuroscience, University College Cork, Ireland; Institute of Environmental Medicine, Karolinska Institutet, Stockholm, Sweden; Department of Psychiatry and Neurobehavioural Science, University College Cork, Ireland

**Author notes:** These Authors contributed equally (John F. Cryan, Lily Keane).

**Keywords:** neurodevelopment, diffuse midline glioma (DMG), microbiota, oligodendrocyte differentiation, microglia, metabolites

## Abstract

Paediatric brain tumours are increasingly recognised as diseases of disrupted development, arising when lineage progression programs, that normally govern neural and glial maturation, become stalled or dysregulated. Diffuse midline glioma (DMG), a highly aggressive paediatric brainstem tumour, emerges during early childhood in the pons, a region undergoing rapid postnatal growth characterized by oligodendrocyte precursor cell (OPC) proliferation and differentiation. However, the environmental factors that shape these developmental trajectories remain poorly defined. Here, using germ-free and conventionally colonized mice, we investigated whether early-life microbiota influences transcriptional and metabolic programs in the developing brainstem during this critical developmental window. Bulk RNA sequencing revealed pronounced microbiota-associated transcriptional differences at postnatal day 2 (P2), but not at P8, identifying a temporally restricted period during which microbial colonization is associated with pathways linked to oligodendrocyte lineage progression, myelination, and neuroimmune signalling. Transcriptional analyses further identified altered expression of genes associated with CD11c⁺ microglia, a developmental microglial subtype implicated in regulating oligodendrocyte maturation. Untargeted metabolomic profiling revealed parallel microbiota-associated differences in pathways related to mitochondrial function, redox balance, and methyl-donor metabolism. Integrated multi-omics analyses identified coordinated networks linking glial lineage programs with metabolites involved in cellular metabolism and epigenetic regulation. Notably, several of these transcriptional and metabolic programs overlap with gene signatures reported in diffuse midline glioma, suggesting that microbiota-sensitive developmental pathways intersect with cellular states relevant to paediatric brainstem tumour biology.

## Introduction

Increasing evidence suggests that many paediatric brain tumours arise from disrupted developmental trajectories in which lineage-specific differentiation programs become stalled or dysregulated(1). Diffuse midline glioma, H3K27-altered (DMG), is classified within the family of paediatric-type diffuse high-grade gliomas (pHGG) and represents an aggressive tumour typically arising in midline brain structures such as the brainstem, thalamus, and spinal cord. The striking spatial and temporal restriction of DMG to the pons and other midline structures strongly suggests that dysregulated developmental programs underlie tumorigenesis (1–4). The pons, the predominant site of DMG, undergoes substantial postnatal expansion driven by proliferation and myelination (7, 8). In humans, pontine volume increases nearly six-fold between birth and five years of age (9), and in mice, a similar burst of growth occurs between postnatal days 0 and 8 (8) . During this period, proliferative Sox2⁺ Olig2⁺ progenitors transition to more differentiated Sox2⁻ Olig2⁺ OPCs by P8 (10), suggesting a lineage progression that, if disrupted, may create a permissive state for tumorigenesis (1, 11).

Cells of the oligodendrocyte lineage are abundant in the developing pons and are thought to represent the cell of origin of DMG (1, 11, 12). The histone H3K27M mutation, characteristic of DMG, locks tumour cells in a progenitor-like, proliferative state and prevents normal differentiation (13, 14). Pharmacologic or genetic restoration of differentiation programs suppresses tumour growth, underscoring the concept that the factors defining postnatal maturation of DMG urgently warrants attention as potential therapeutic targets (13, 14). Identifying factors that influence oligodendrocyte precursor cell (OPC) proliferation and differentiation during this vulnerable developmental period is therefore critical for understanding disease susceptibility.

Microglia, the immune cells of the brain, are highly dynamic, heterogeneous and are essential regulators of postnatal brain development (13, 14). They maintain neuronal health and activity, sculpt neuronal circuits through synaptic pruning (17–19), and regulate OPC proliferation, differentiation, and myelination (20–22). Microglial are also incredibly diverse with temporal and regionally distinct microglia subtypes previously described (23–29). DMG exhibits a uniquely immunologically cold tumour microenvironment that contributes to disease progression(28, 29). The immune landscape of DMG differs markedly from that of adult-type diffuse gliomas such as glioblastoma. Comparative studies using immunocompetent de novo mouse models of paediatric high-grade glioma have demonstrated that both the underlying driver mutations and the anatomical site of origin shape the tumour microenvironment (30). Importantly, DMG tumours have been shown to have more affinity for growth and invasion in the brainstem compared to cortical regions of the brain during early postnatal development (31).

The gut microbiota has recently emerged as a key regulator of brain development, with coordinated postnatal assembly shaping neurodevelopmental processes including microbial maturation and function (33–35). Bidirectional communication between the gut microbiota and brain, also known as the microbiota-gut-brain axis, is mediated through immune, endocrine, and metabolic pathways (44, 45). During the first three years of life, microbial colonization dominated by *Bifidobacterium* and *Lactobacillus* species also establishes a foundation for neuroimmune and metabolic signalling (36, 37) and perturbations of the microbiota during this period have been linked to neurodevelopmental disorders (38). Microbiota-derived metabolites—including short-chain fatty acids (SCFAs), tryptophan derivatives, and neurotransmitters such as serotonin and γ-aminobutyric acid (GABA) modulate brain activity and influence key functions such as blood–brain barrier integrity (39), myelination (40), neurogenesis (41), and microglial maturation and function (42, 43). Indeed, microbial metabolites, are essential for microglial maturation and function through epigenetic mechanisms involving histone methylation/acetylation (42, 43, 47). Importantly previous work from our lab has shown that disrupting the early life microbiota at critical windows during development in the prefrontal cortex, can have lasting effects on microglia, behaviour and myelination(34, 46). However, it is still unclear how microbial signals molecules reprogram microglial subtypes, or how gut microbiota–microglia interactions influence oligodendrocyte lineage progression and brainstem maturation during early postnatal development.

To this end, using germ-free and conventionally raised mice, we characterise microbiota-dependent transcriptional and metabolic programs in the developing brainstem that are associated with microglial activation and oligodendrocyte lineage progression. Our goal is to define a microbiota-sensitive developmental framework linking early-life environmental cues to brainstem maturation, and to propose a potential developmental context for vulnerability to diffuse midline glioma, while providing a resource for future mechanistic and disease-focused studies.

## Results

### Microbiota-dependent transcriptional differences are most pronounced at postnatal day 2

To determine whether microbiota-dependent transcriptional programs differ across early postnatal stages of brainstem development, we focused on two time points that bracket rapid pons growth and subsequent stabilization(7, 8) i.e. postnatal day 2 (P2) or postnatal day 8 (P8) germ-free (GF) and conventionally raised (CONV) male mice and performed RNA sequencing (Figure 1A). P2 was selected as it corresponds to the critical period of postnatal pons expansion in the mouse brain (4-times increase between P0-P4) whereas P8 represents a more static period of postnatal growth (7, 8). Principal component analysis (PCA) revealed clear separation between GF and CONV samples at P2 but not at P8, indicating that microbial status exerts its strongest transcriptional effect during the earliest phase of postnatal development (PERMANOVA - comparison between germfree versus conventional, hereafter referred to as microbiota status,: P2 (F = 3.04, R² = 0.178, p = 0.029); P8 (F = 1.11, R² = 0.073, p = 0.317)) (Figure 1B). Permutational multivariate analysis of variance (PERMANOVA) confirmed a trend for significant interaction between age and microbiota status (F = 1.53, R² = 0.141, p = 0.1), consistent with a time-restricted transcriptional response. Interestingly, brainstem transcriptome analysis also identified an influence of age in GF but not in CONV mice (PERMANOVA - age: GF (F = 2.34, R² = 0.143, p = 0.039); CONV (F = 1.41, R² = 0.091, p = 0.22)) (Supplemental Figure 1A-1C). Differential expression analysis identified 4,712 genes significantly altered (q < 0.2) between GF and CONV brainstems at P2, compared to no genes being differentially expressed at P8 (Figure 1C and Supplemental Figure 1C). The limited overlap between these gene sets suggests that the potential effect of the microbiota on brainstem transcriptional programs is temporally restricted. Volcano plots highlighted robust up-and down-regulation of genes associated with cell cycle regulation, myelination, and immune signalling in GF compared to CONV mice at P2, with no significant differences being observed at P8 (Figure 1D). Top differentially expressed genes further demonstrated transcriptional divergence between GF and CONV mice was most pronounced at P2, with several microglia-and oligodendrocyte-related transcripts markedly altered in GF brainstems (Figure 1E). Together, these results indicate that P2 represents a period of heightened microbiota sensitivity, during which microbial status is associated with pronounced transcriptional differences in the developing brainstem.

**Figure 1.**
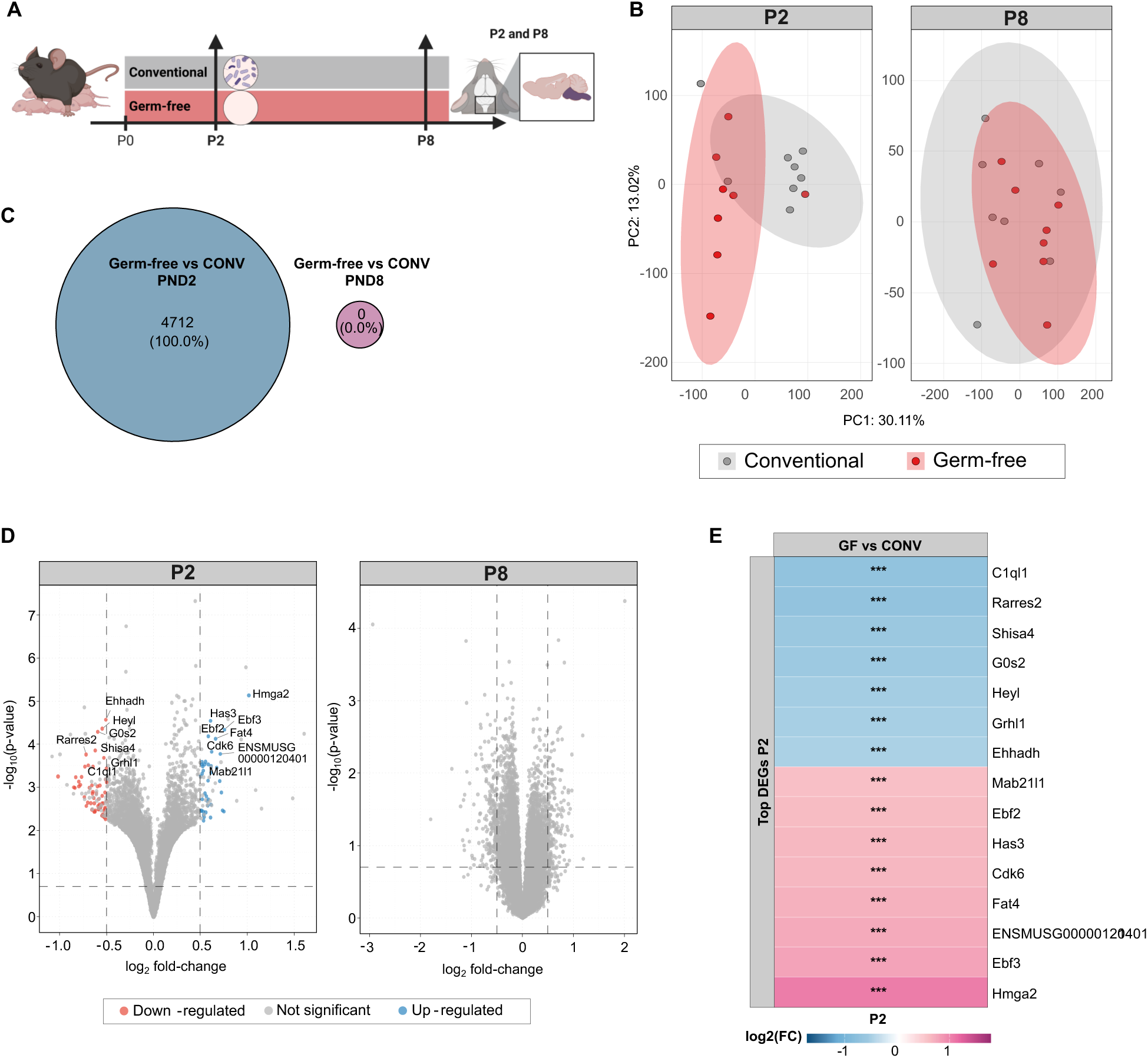
Postnatal day 2 marks a critical window for microbiota-driven transcriptional programs in the developing brainstem. **A**) Schematic representation of the experimental design for RNAseq in the brainstem of germ-free and conventional mice at post-natal day 2 (P2) and P8 (n = 8 mice/group/age). **B**) Principal Component Analysis (PCA) plot showing differences in the brainstem transcriptome of microbiota status (germ-free vs conventional) mice at P2 and P8. Data analysed using PERMANOVA analysis. **C**) Venn diagram showing overlap of significantly DEGs (*q < 0.2*) between germ-free and conventional mice at P2 and P8. **D**) Volcano plot illustrating the most differentially expressed genes (DEGs) (P-adjusted (*q*) value<0.2 and log₂-fold change≥ 0.5; up-regulated (blue) and down-regulated (red) genes, with the top 15 DEGs for germ-free vs conventional mice at P2 and P8) in the brainstem. **E**) Heatmap showing differential expression (log_2 f_old change) of the top most DEGs in the brainstem of germ-free vs conventional mice at P2 and P8 (colours represent direction and magnitude of expression change (purple = upregulated in germ-free; blue = downregulated), and asterisks indicate statistical significance of P-value (*\*p<0.05, **p<0.01, ***p<0.001*) for microbiota status).

### Early-life microbiota modulates oligodendrocyte lineage–associated transcriptional programs intersecting with diffuse midline glioma gene signatures

Having established that P2 represents a microbiota-sensitive transcriptional window in the developing brainstem, we next wanted to examine how the absence of the microbiota could affect oligodendrocyte lineage progression. Oligodendrocyte lineage development proceeds through a tightly regulated sequence from proliferative pre-OPCs to mature myelinating oligodendrocytes (47). We therefore carried out differential expression analysis on sets of genes relating to key stages of oligodendrocyte differentiation and found downregulation of genes across all stages of oligodendrocyte differentiation in GF compared to CONV mice at P2 but not at P8 (Figure 2A). This suggests that transcriptional programs controlling early progenitor specification through myelin gene expression, are suppressed in germ-free mice and that microbial cues are required continuously throughout the oligodendrocyte differentiation trajectory to drive proper oligodendrocyte maturation and myelination.

**Figure 2.**
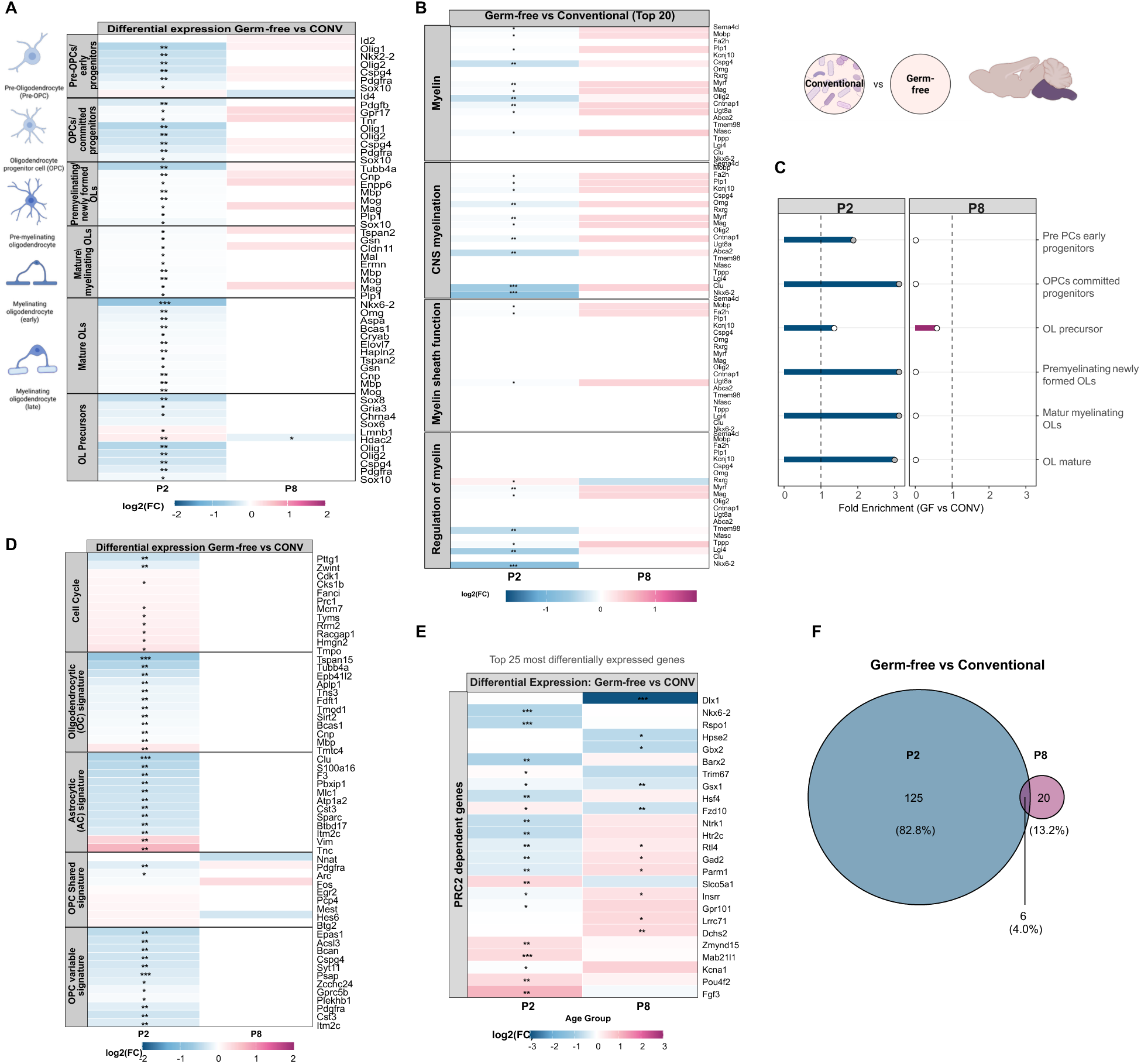
Early life microbiota is critical for oligodendrocyte precursor cell (OPC) proliferation and differentiation in the developing brainstem. **A-B**) Heatmaps showing differential expression (log2 fold change) of top genes associated with A) oligodendrocyte differentiation, **B**) myelin regulation, myelination in the central nervous system (CNS), myelin function and myelin regulation in germ-free vs conventional mice at post-natal day 2 (P2) and P8. **C**) Targeted enrichment analysis of manually curated gene sets associated with oligodendrocyte and myelin-related pathways in germ-free versus conventional male mice at P2 and P8. Analysis performed on significantly differentially expressed genes (*p<0.05*). Bars represent fold enrichment values, with grey circles indicating significant enrichment (*p<0.05*) and white circles indicating non-significant enrichment. Dashed line represents no enrichment (fold change = 1). D-E) Heatmap showing differential expression (log2 fold change) of the top DEGs associated with **D**) H3K27M glioma signatures and the **E**) 25 most DEGs associated with Polycomb Repressive Complex 2 (PRC2) in the brainstem of germ-free vs conventional mice at P2 and P8. **F**) Venn diagram showing overlap of significantly differentially expressed PRC2-dependent genes (*p<0.05*) between germ-free and conventional mice at P2 and P8. A-B, D-E) Colours represent direction and magnitude of expression change (purple = upregulated in germ-free; blue = downregulated), and asterisks indicate statistical significance of P-value (\**p<0.05*, \*\**p<0.01*, \*\*\**p<0.001*) for microbiota status) (n = 8 mice/group/age).

Indeed, differential gene expression analysis also revealed widespread dysregulation of genes involved in myelin regulation and formation in GF mice compared to CONV mice at P2, but not at P8 (Figure 2B). To assess the broader impact on oligodendrocyte-related pathways, we next performed targeted enrichment analysis of differentially expressed genes (p < 0.05). Gene sets linked to OPC proliferation, differentiation, and myelination were significantly enriched among genes decreased in GF mice at P2, further confirming that microbiota absence delays oligodendrocyte maturation (Figure 2C and Supplemental Figure 1D).

Diffuse midline glioma (DMG), H3K27M mutant, arises within the brainstem during early childhood, and its striking spatial confinement to the pons and temporal restriction to early postnatal development suggest that it originates from stalled or dysregulated neurodevelopmental programs (1). Recently, Filbin et al., showed that H3K27M gliomas contain cells resembling OPCs (OPC-like), which displayed higher proliferation and tumour-invasion potential, and they identified low number of differentiated tumour cell (10).

Comparison with published H3K27M-associated gene signatures revealed increased overlap with DMG-related transcriptional programs in GF brainstems at P2 (Figure 2D). These signatures were enriched for genes associated with progenitor-like and cell-cycle–related processes, consistent with a less differentiated transcriptional profile.

Because H3K27M mutation–driven gliomas are characterized by epigenetic dysregulation of Polycomb Repressive Complex 2 (PRC2)–dependent genes, as defined by a previous study (48), we next examined whether these programs were similarly altered in GF mice during early postnatal development. Heatmap visualization revealed differential expression of multiple PRC2-regulated targets, with a predominance of upregulation in GF mice (Figure 2E). Germ-free mice had a larger number of PRC2-dependent genes differentially expressed compared to CONV mice at P2 than at P8 (125 vs 20, respectively), again underscoring the temporal specificity of microbiota influence (Figure 2F). Additionally, a significant enrichment in pathways related to histone modifiers and metabolic regulation as well as node of Ranvier, juxtaparanode, paranode and internode was observed in GF vs CONV mice at P2, and in GABA transmission at P8 (Supplemental Figure 1E and F).

Together, these data suggest that early-life microbiota status influences transcriptional programs linked to OPC differentiation and myelination, potentially through metabolically mediated epigenetic mechanisms, and that its absence is associated with progenitor-related transcriptional signatures overlapping with those reported in diffuse midline glioma.

### Early-life microbiota modulates microglia-associated transcriptional programs linked to CD11c⁺ signatures, a developmental population enriched in diffuse midline glioma

We next investigated whether these effects extend to microglial populations known to regulate OPC maturation and are themselves implicated in DMG biology. Given the convergence between microbiota-dependent transcriptional programs and DMG-like gene signatures, we next examined whether specific microglial populations implicated in brainstem development and gliomagenesis were affected by microbial status (Figure 3A–C). Heatmap analysis of published brainstem microglial signatures revealed that CD11c⁺ microglia are the predominant microglia population present in DMG patients (Figure 3A and Supplemental Figure 2A). Interestingly, we found, for the first time to the best of our knowledge, that these CD11c⁺ microglia gene signatures were also dysregulated in germ-free mice compared to CONV controls at P2 (Figure 3B). In contrast, signatures associated with other microglia subtypes including keratan sulfate proteoglycan (KS⁺), Hoxb8⁺, and Arg1⁺ microglia were similar between germ-free and CONV controls at both P2 and P8 (Supplemental Figure 2B).

**Figure 3.**
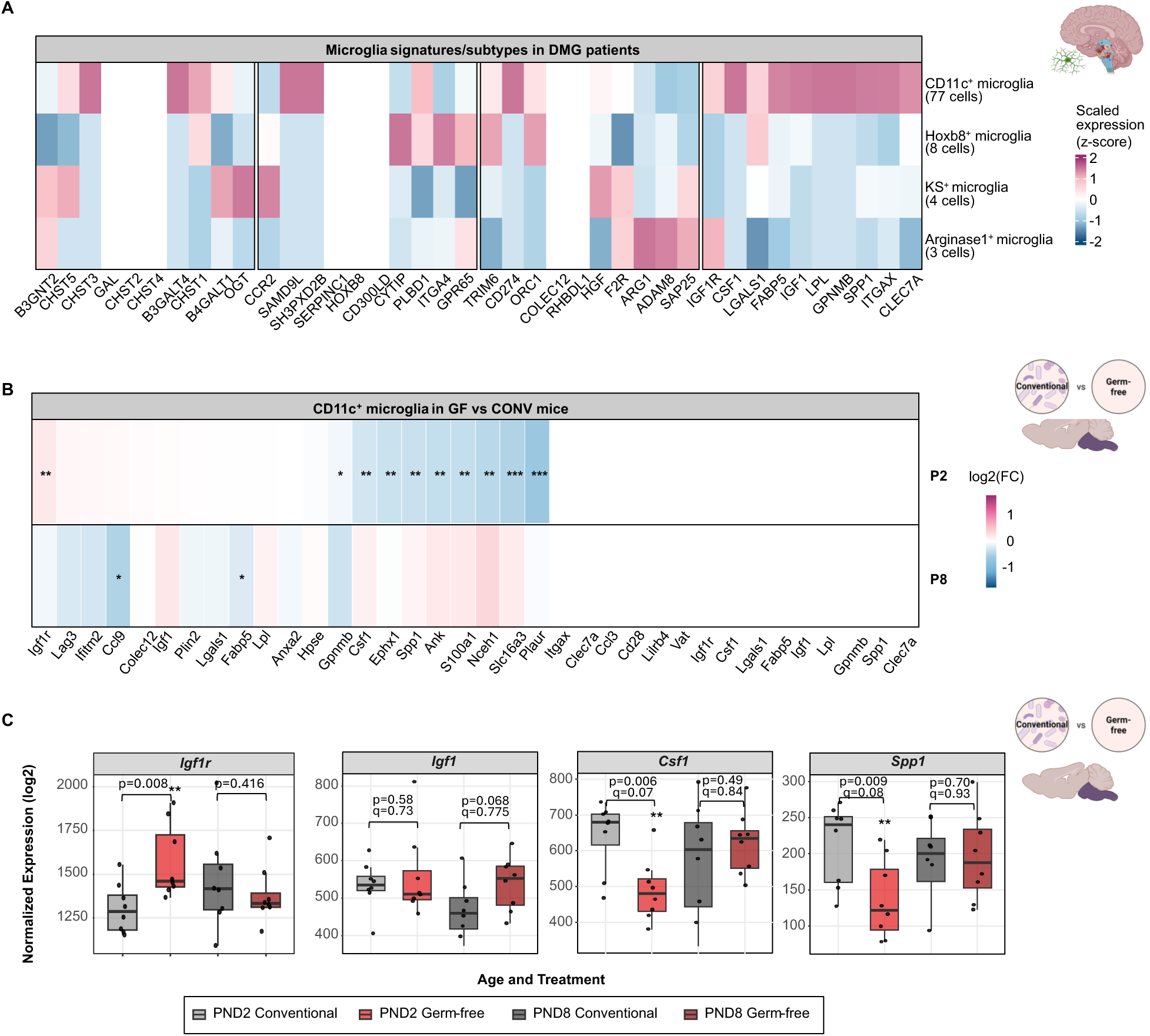
CD11c^+^ microglia dominant in DMG tumours and are dysregulated in the brainstem of germ-free mice. **A**) Heatmap showing scaled average expression (z-scores) of signature genes associated with four distinct microglia subtypes (KS^+^, Hoxb8^+^, Arg1^+^, and CD11c^+^) present in H3K27M mutant diffuse midline gliomas (DMG). Data are derived from brainstem single-cell RNA-seq of 92 immune cells from human DMG patient tissue. **B**) Heatmap showing differential expression (log₂ fold change) of CD11c^+^ microglia gene signatures in the brainstem of germ-free versus conventional mice at postnatal day 2 (P2) and P8. Colours represent direction and magnitude of expression change (purple = upregulated in germ-free; blue = downregulated). **C**) Boxplots depicting normalized expression (log₂) of *Igf1r, Igf1, Csf1* and *Spp1* in brainstem of germ-free and conventional mice at P2 and P8. Data are shown as median ± IQR, with individual points representing biological replicates. Differential expression results (log₂FC, P-adj (*q*)) were calculated with DESeq2. **B-C**) Asterisks indicate statistical significance of P-value (\**p<0.05*, \*\**p<0.01*, \*\*\**p<0.001*) for microbiota status) (n = 8 mice/group/age).

At postnatal day 2 (P2), numerous CD11c⁺-associated transcripts, including *Igf1*, Csf1, *Spp1*, and *Ank1* were significantly downregulated in GF brainstems, while few differences persisted by P8 (Figure 3B–C and Supplemental Figure 2C). These genes encode proteins essential for CD11c+ microglial activation, metabolic reprogramming, and trophic support of oligodendrocyte precursor cells (49), suggesting that absence of microbial-derived signals, such as metabolites, disrupts the emergence or maintenance of this specialized microglial subset during a critical developmental window. Together, these findings indicate that early-life microbiota status modulates microglia-associated transcriptional programs overlapping with CD11c⁺ signatures reported in diffuse midline glioma, highlighting a potential connection between microbial cues and neuroimmune maturation during brainstem development

### Early-life microbiota shapes brainstem metabolic profiles during early postnatal development

Given that microbial signals can modulate microglial maturation through metabolite-mediated pathways, we next performed untargeted metabolomic profiling of germ-free and conventional brainstems at P2 and P8 (Figure 4A–D), to identify potential metabolic cues underlying microbiota-dependent regulation of brainstem development. Principal component analysis revealed clear segregation in metabolite profiles between GF and CONV samples at P2 but not at P8, indicating that microbial colonization exerts its strongest metabolic influence during the earliest phase of postnatal life (PERMANOVA -microbiota status: P2 (Pseudo F = 4.53, R² = 0.312, p = 0.003); P8 (Pseudo F = 0.55, R² = 0.052, p = 0.733); age: GF (Pseudo F = 10.65, R² = 0.518, p = 0.002); microbiota status (Pseudo F = 8.25, R² = 0.452, p = 0.003)) (Figure 4A). Indeed, differential abundance analysis identified 34 significantly altered metabolites in GF versus CONV brainstems at P2 (p < 0.05, log₂ fold change ≥ 0.3), with markedly fewer differences by P8 (Figure 4B). Our results showed that most differentially abundant metabolites were reduced in GF mice, suggesting depletion of microbiota-derived or microbiota-modulated compounds (Figure 4C).

**Figure 4.**
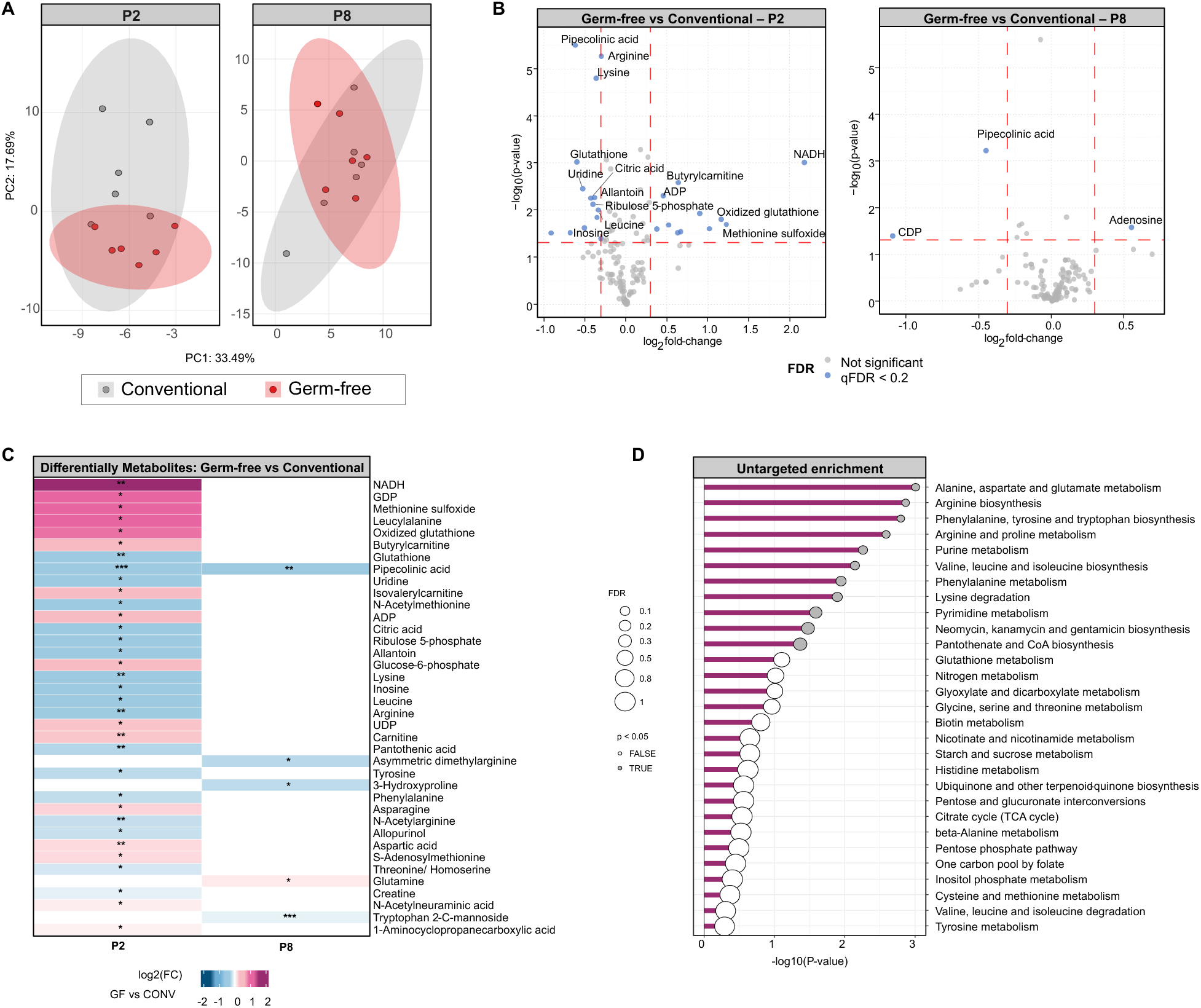
Microbial metabolites are altered in germ-free mice at postnatal day 2, a critical window in brainstem development. **A**) Principal Component Analysis (PCA) plot showing differences in the brainstem metabolome of germ-free and conventional mice at postnatal day 2 (P2) and P8 (n = 8 mice/group/age). **B**) Volcano plot illustrating differentially abundant metabolites (p < 0.05 and log₂-fold change ≥ 0.3) in germ-free vs conventional mice at P2 and P8. Blue points indicate significantly differentially abundant metabolites, with the top 15 most significant metabolites labelled for each timepoint. **C**) Heatmap showing the differentially abundant metabolites (log2 fold change, germ-free vs conventional) in the brainstem at P2 and P8 (colours represent direction and magnitude of abundance changes (blue = decreased in germ-free; red = increased in germ-free), and asterisks indicate statistical significance of P-value (*\*p<0.05, **p<0.01, ***p<0.001*) for microbiota status). **D**) Untargeted enrichment analysis of metabolites that were significantly altered in germ-free compared to control mice at P2 (*p<0.05*).

Among the metabolites most decreased in GF mice at P2 were pipecolinic acid, glutathione, and citric acid, key regulators of immune modulation, mitochondrial function and oxidative stress. In contrast, metabolites that were more increased were NADH, oxidized glutathione and GDP implicating cellular redox homeostasis and energy metabolism (Figure 4B and 4C). Interestingly, S-adenosylmethionine, a key metabolite for epigenetic methylation and previously implicated in DMG pathogenesis (50), was also significantly increased in GF compared to CONV mice at P2 (Figure 4C). Indeed, recent work demonstrates that methionine cycle activity and SAM flux are elevated in DMG, and that methionine or SAM restriction suppresses tumour growth by modulating PRC2-dependent methylation (50). Untargeted pathway enrichment analysis revealed significant perturbation of lysine degradation, arginine biosynthesis, and other amino acid metabolic pathways (Figure 4D), consistent with reduced metabolic support for neurodevelopmental and glial processes. Together, these findings support a role for early-life microbiota in shaping the metabolic landscape of the developing brainstem during early postnatal life, intersecting with pathways relevant to microglial and oligodendrocyte maturation.

### Integrated metabolic and transcriptional pathways associated with microglia and OPC programs in the early postnatal brainstem

To integrate microbiota-dependent changes across molecular layers, we performed an untargeted pathway analysis combining differentially expressed genes and metabolites from GF and CONV brainstems at P2, which corresponds to the critical period of postnatal pons expansion and was identified as the key window for the strongest microbiota influence on brainstem development (Figure 5A). Integrated analysis using MetaboAnalyst 6.0 revealed strong convergence of transcriptional and metabolic alterations within pathways central to cellular energy metabolism, amino-acid biosynthesis, and epigenetic regulation (Figure 5A).

**Figure 5.**
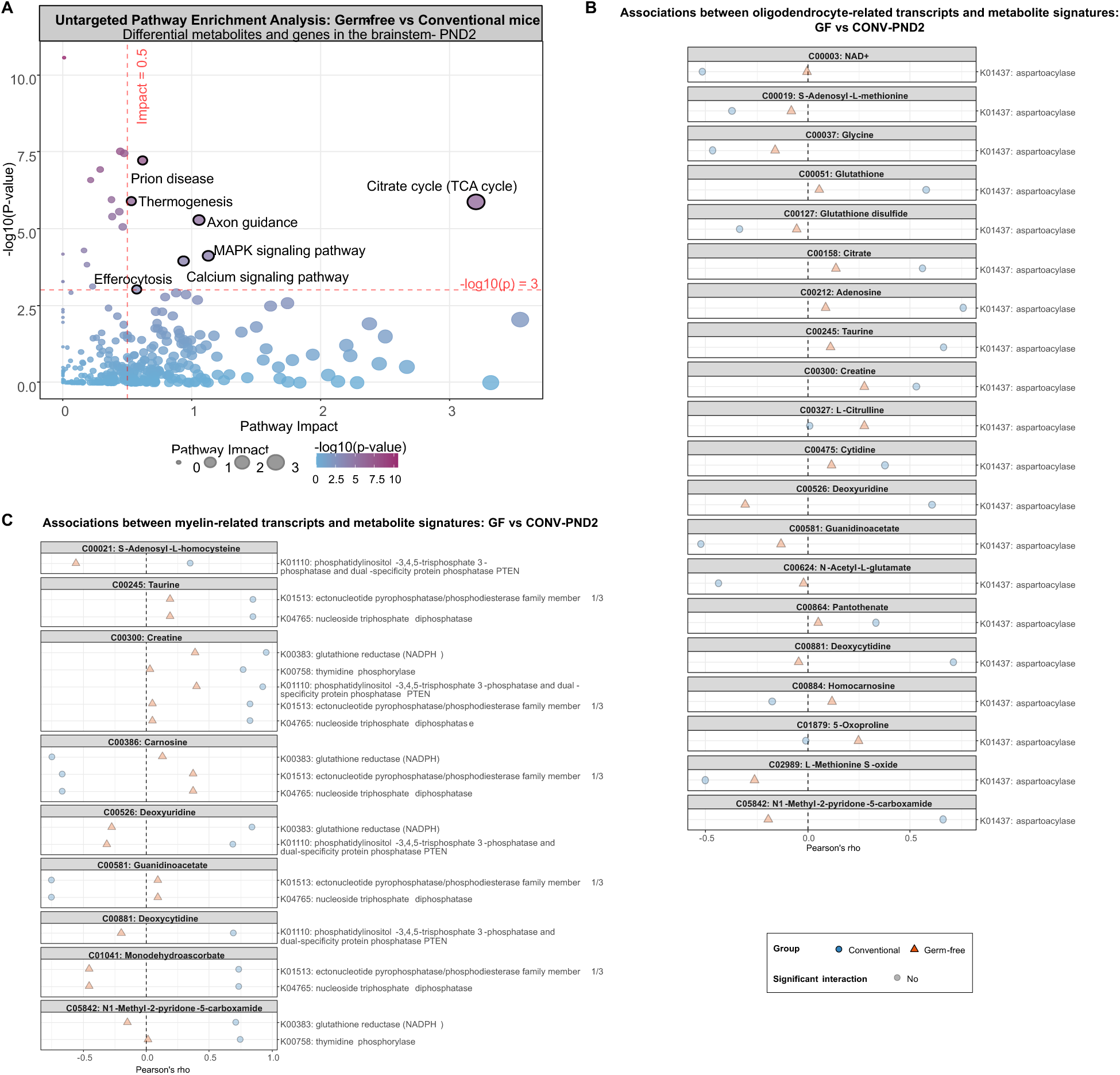
Multi-omics integration identifies microbiota-dependent metabolic networks linked to oligodendrocyte maturation and myelination at P2. **A**) Untargeted integrated pathway analysis in brainstem samples of germ-free vs conventional mice at postnatal day 2 (P2). Analysis based on differentially expressed genes (RNA-seq, p-value<0.05) and differentially abundant metabolites (metabolomics, p-value<0.05) using MetaboAnalyst 6.0 (n = 8 mice/group). Circle size reflects pathway impact; colour intensity represents statistical significance (light blue to purple). Pathways with high biological impact (≥0.5) and statistical significance (-log10(p) ≥ 3; p < 0.001) are highlighted with black borders and labelled. **B-C**) Correlation between **B**) oligodendrocyte differentiation-and **C**) myelin-related transcripts and metabolite signatures sharing a KEGG pathway at P2. Data points represent Pearson correlation coefficients calculated between corresponding (**B**) oligodendrocyte differentiation-and **C**) myelin-related) gene expression and metabolite abundances in conventional (circles) and germ-free (triangles) groups. Opaque symbols denote statistically significant differential correlations between microbiota states (Benjamini-Hochberg corrected *q<0.05* for between-group comparison), whereas translucent symbols represent non-significant differences. Gene sets examined included **B**) stage-specific markers of oligodendrocyte maturation (pre-OPCs, committed OPCs, premyelinating oligodendrocytes, and mature myelinating oligodendrocytes) and **C**) CNS myelination, myelin sheath function, myelination, regulation of myelin and myelin structural proteins.

Next, we performed joint-pathway analysis for microbiota influence on oligodendrocyte-related metabolic networks. Integrated analysis showed that GF mice at P2 had lower or altered associations for metabolites related to mitochondrial function (such as NAD⁺, citrate, creatine), redox regulation (glutathione, carnosine), and methylation (such as S-adenosyl-L-methionine, L-methionine S-oxide) (Figure 5B). Additionally, GF mice displayed lower associations between myelin-related transcripts and metabolites key for mitochondrial energy metabolism (including isocitrate, creatine/creatinine coupling), antioxidant (e.g. carnosine), and epigenetic regulation (e.g. S-adenosyl-L-methionine) at P2 (Figure 5C). GF mice at P2 also exhibited reduced associations between PRC2 complex–related transcripts and metabolites involved in methylation and nucleotide metabolism, such as creatine or deoxyuridine, linked to cGMP signalling, lipid metabolism and extracellular matrix synthesis and cell-surface glycosylation (Figure 6A). Further analysis revealed reduced associations between H3K27M-associated transcripts and metabolites linked to mitochondrial, nucleotide and methylation-linked metabolism (including creatine, carnitine, deoxyuridine, deoxycytidine) at P2 (Figure 6B). At P2, GF mice had lower correlations in KOs associated with oxidative phosphorylation and electron transport, pyrimidine and purine biosynthesis and acyl-carnitine metabolism (Figure 6B). Lastly, we performed integrative analysis for CD11c^+^ microglial signature which showed stronger correlations with mitochondrial redox and nucleotide turnover metabolites (e.g. butyrylcarnitine, NADH, CDP, GDP, UDP), and reduced correlations with glutathione, glycolytic and methyl donor metabolites (e.g. glutathione, lactic acid, nicotinamide) in GF mice at P2 (Figure 6C).

**Figure 6.**
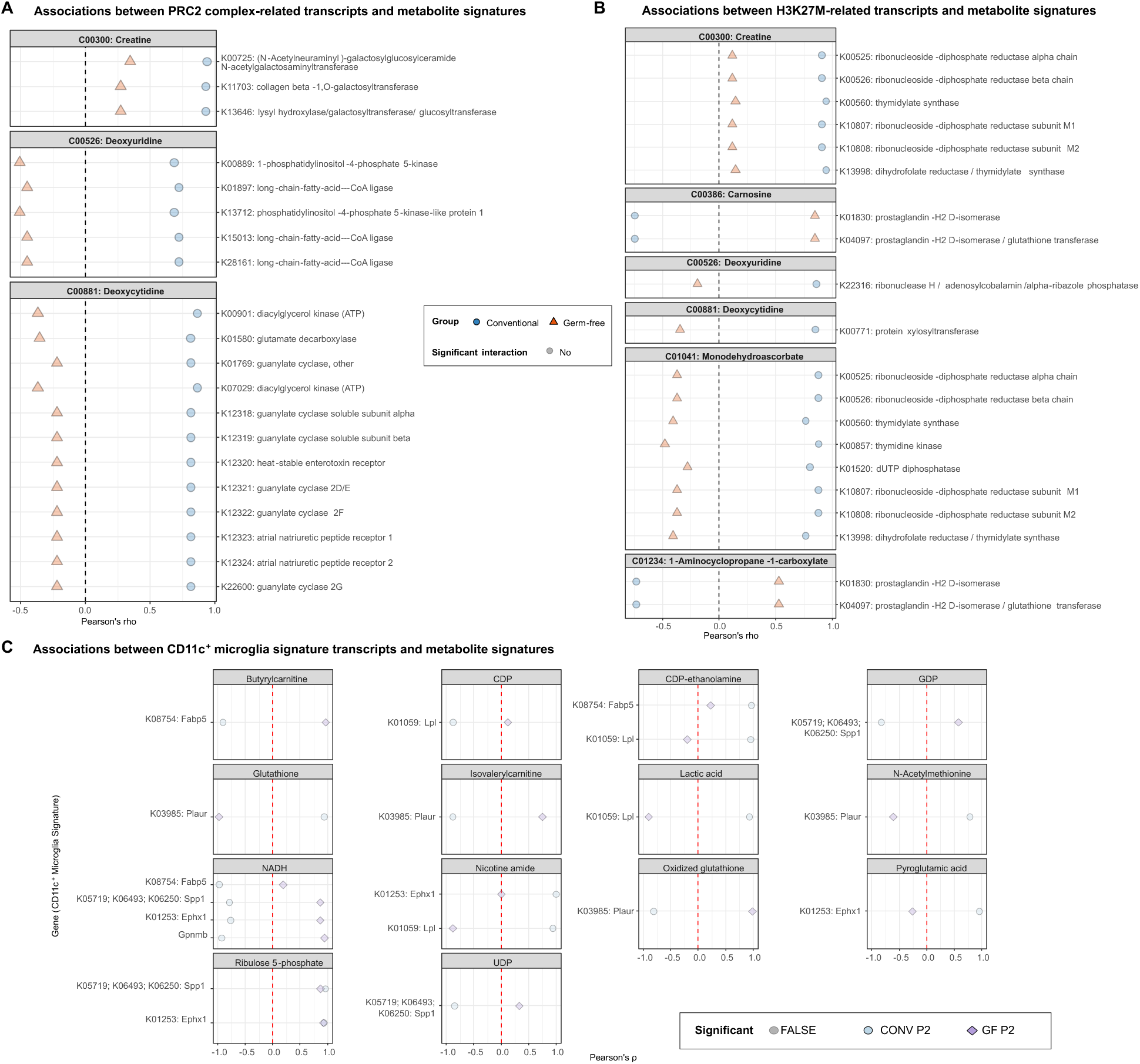
Multi-omics integration identifies microbiota-dependent metabolic networks linked to PRC2 complex and H3K27M mutation at P2. **A-C**) Correlation between **A**) PRC2 complex- and B) H3K27M-related transcripts and metabolite signatures sharing a KEGG pathway at P2. Data points represent Pearson correlation coefficients calculated between corresponding (**A**) PRC2 complex-and **B**) H3K27M-related) gene expression and metabolite abundances in conventional (circles) and germ-free (triangles) groups. Opaque symbols denote statistically significant differential correlations between microbiota states (Benjamini-Hochberg corrected q<0.05 for between-group comparison), whereas translucent symbols represent non-significant differences. Gene sets examined included stage-specific markers of **A**) PRC2 complex and **B**) cell cycle signatures-H3K27M Glioma, OC signatures-H3K27M Glioma, AC signatures-H3K27M Glioma, OPC shared signatures-H3K27M Glioma and OPC variable signatures-H3K27M Glioma. **C**) Correlation between CD11c^+^ microglia signature transcripts and metabolites in P2. Data points represent Pearson correlation coefficients (ρ) between gene expression and metabolite abundance for conventional (blue circles) and germ-free (purple diamonds) mice. Filled symbols indicate significant differential associations between microbiota states (Benjamini-Hochberg adjusted interaction *q<0.05*), while transparent symbols indicate non-significant interactions. Only associations with overall correlation Benjamini-Hochberg adjusted *q<0.1* and interaction *p<0.05* are shown. Gene labels include KEGG Orthology IDs where available.

Several of these pathways directly intersect with the metabolites depleted in GF brainstems at P2, including carnitine, glutathione, and methyl-donor intermediates, suggesting that microbial-derived or microbially regulated metabolites provide key substrates for OPC differentiation or microglia activation. This integrated analyses reveal coordinated microbiota-associated metabolic and transcriptional networks in the early postnatal brainstem that intersect with microglia-and OPC-associated programs, suggesting a potential connection between microbial metabolism and pathways relevant to diffuse midline glioma.

Collectively, our findings support the existence of an early postnatal period during which microbiota-associated metabolic, immune, and glial programs are co-ordinately altered in the developing brainstem, providing a foundation for therapeutic strategies that target microbiota-derived metabolites to modulate neurodevelopment and DMG vulnerability

## Discussion

The present study highlights early-life gut microbiota status as an important modifier of postnatal brainstem development, associated with coordinated transcriptional, metabolic, and neuroimmune programs. By integrating RNA sequencing and metabolomic analyses from germ-free and conventionally raised mice, we identify postnatal day 2 (P2) as an early postnatal stage at which microbiota-associated differences in brainstem gene expression and metabolic profiles are most pronounced. At this stage, microbial status was associated with transcriptional programs linked to oligodendrocyte lineage progression, myelination, and microglia-related pathways.

Absence of the microbiota during early postnatal development coincided with widespread transcriptional and metabolic alterations in the brainstem, including enrichment of progenitor-and cell-cycle–associated gene signatures that overlap with those reported in diffuse midline glioma (DMG). While these findings do not imply disease causation, they suggest that microbiota-sensitive developmental programs intersect with cellular states relevant to DMG biology. Our data therefore supports the existence of a microbiota-sensitive early postnatal period during which metabolic and neuroimmune pathways associated with microglia and oligodendrocyte maturation are dynamically regulated, providing a developmental context for exploring how early-life microbial cues may influence brainstem vulnerability to disease.

### OPC differentiation, H3K27M/PRC2 biology, and SAM metabolism

Paediatric brain tumours such as DMG are increasingly recognized as disorders of dysregulated neurodevelopment, arising from progenitor populations confined within spatially and temporally restricted niches (1, 3). The localization of DMG to the pons during early childhood coincides with a developmental window dominated by Olig2⁺ OPC proliferation and differentiation (7, 8). Single-cell and multi-omic studies reveal that DMG tumour cells predominantly adopt an OPC-like identity and remain stalled along a restricted differentiation trajectory (9). Experimentally, expression of H3K27M in neural progenitors reprograms cells toward a primitive, self-renewing state, whereas its removal restores H3K27me3 deposition, leading to the reactivation of glial differentiation programs, and the attenuation of tumour growth (51, 52). Mechanistically, H3K27M retargets PRC2 complex activity, maintaining repression at lineage-defining loci while sustaining enhancer activity at progenitor-associated genes (53).

Our findings extend this developmental framework by suggesting that the early-life microbiota may provide an additional layer of regulation over these intrinsic lineage programs that warrants further investigation. In germ-free mice, OPCs revealed transcriptional programs suggesting stalled differentiation, with downregulation of *Myrf, Sox10, Mbp, and Plp1* genes, and upregulation of progenitor and cell-cycle transcripts, including *Cks1b, Hmgn2, Mcm7, Racgap1, Rrm2, Tmpo* and *Tyms*, mirroring the transcriptional and epigenetic immaturity of H3K27M DMG tumour cells. Dysregulation of key microbial metabolites such as S-adenosylmethionine indicates a metabolic– epigenetic mechanism, whereby microbial cues stabilize PRC2 activity and promote OPC maturation. Indeed, S-adenosylmethionine (SAM) is the universal methyl donor for histone methyltransferases, including EZH2, the catalytic subunit of the Polycomb Repressive Complex 2 (PRC2)(54). EZH2 catalyses the conversion of H3K27 to H3K27me3 only when SAM is available, producing S-adenosylhomocysteine (SAH), which in turn inhibits EZH2 if accumulated (55). Accordingly, both deficiency and imbalance of SAM can alter global methylation and disrupt H3K27me3 homeostasis. PRC2 activity is highly sensitive to one-carbon metabolism, which maintains SAM through the folate and methionine cycles (56). In H3K27M DMG, the oncohistone binds and inhibits PRC2, leading to loss of H3K27me3 globally but aberrant maintenance of repression at developmental genes, locking cells in an undifferentiated, progenitor-like state (1, 53). Importantly, recent work demonstrates that methionine cycle activity and SAM flux are elevated in DMG, and that methionine or SAM restriction suppresses tumour growth by modulating PRC2-dependent methylation (50). In our study, SAM levels were increased in germ-free brainstems at postnatal day 2, the stage showing OPC transcriptional programs reflecting stalled differentiation as well as PRC2 target gene dysregulation, suggesting that microbial signals normally buffer methyl-donor availability. The absence of these cues may hyperactivate or mislocalize PRC2, reinforcing repressive chromatin at differentiation loci.

Our data also showed that microbiota-dependent alterations in aspartoacylase (K01437)-linked metabolism were strongly correlated with the expression of myelin-associated transcripts in the early postnatal brainstem. Aspartoacylase catalyzes the hydrolysis of N-acetylaspartate (NAA) to generate aspartate and acetate, a critical substrate for lipid biosynthesis during myelination (57). The identification of this pathway among the top-ranked microbiota-sensitive nodes suggests that gut microbial signals influence central acetate metabolism, potentially shaping myelin formation during early development.

### Microglia-OPC interactions

Microglia are key developmental regulators of oligodendrocyte lineage progression, influencing both OPC proliferation and differentiation through direct contact and trophic signalling, including IGF1, SPP1, CSF1 and ANK1. A specialized population of CD11c⁺ microglia, also termed proliferative region–associated or axon tract–associated microglia, emerges transiently within white matter tracts during early postnatal development (P2–P7 in mice), coinciding with the peak of OPC expansion and myelination (49, 58). These cells express high levels of IGF1 and SPP1, which act on nearby OPCs to promote differentiation and myelin sheath formation (59). In their absence, developmental myelination is markedly impaired, demonstrating that CD11c⁺ microglia are essential for establishing proper white matter architecture (49). Microglia also contribute to myelin refinement through phagocytic remodelling of redundant sheaths (19), and disruption of microglial homeostasis, such as in FireΔ/Δ mice lacking microglia, results in early hypermyelination followed by demyelination, underscoring their role in maintaining balanced myelin turnover (20). Microglial maturation is metabolically and epigenetically tuned by microbiota-derived SCFAs, which regulate chromatin accessibility and activation(14, 43, 44).

Analysis of single-cell transcriptomic datasets from human DMG revealed a predominance of CD11c⁺ microglia suggesting DMG co-opts a developmental microglial program that supports progenitor proliferation while maintaining immune suppression. Consistent with this, our previous work showed that deletion of EZH2 epigenetically reprograms microglia toward an anti-tumoural activation state, underscoring the plasticity of these cells (60). Collectively, these findings imply that the same microglial subtype essential for early myelination may be re-engaged in DMG to sustain tumour growth and immunosuppression. In our study, germ-free mice exhibited transcriptional dysregulation of CD11c⁺ microglial markers (Igf1r, Spp1, Csf1) and impaired OPC differentiation (reduced Myrf, Sox10, Mbp, and Plp1), mirroring the phenotype observed when microglial trophic support fails. Together, these findings support a role for microbiota-dependent microglial signalling in shaping transcriptional programs associated with OPC maturation and myelination during early brainstem development (Figure 7).

**Figure 7.**
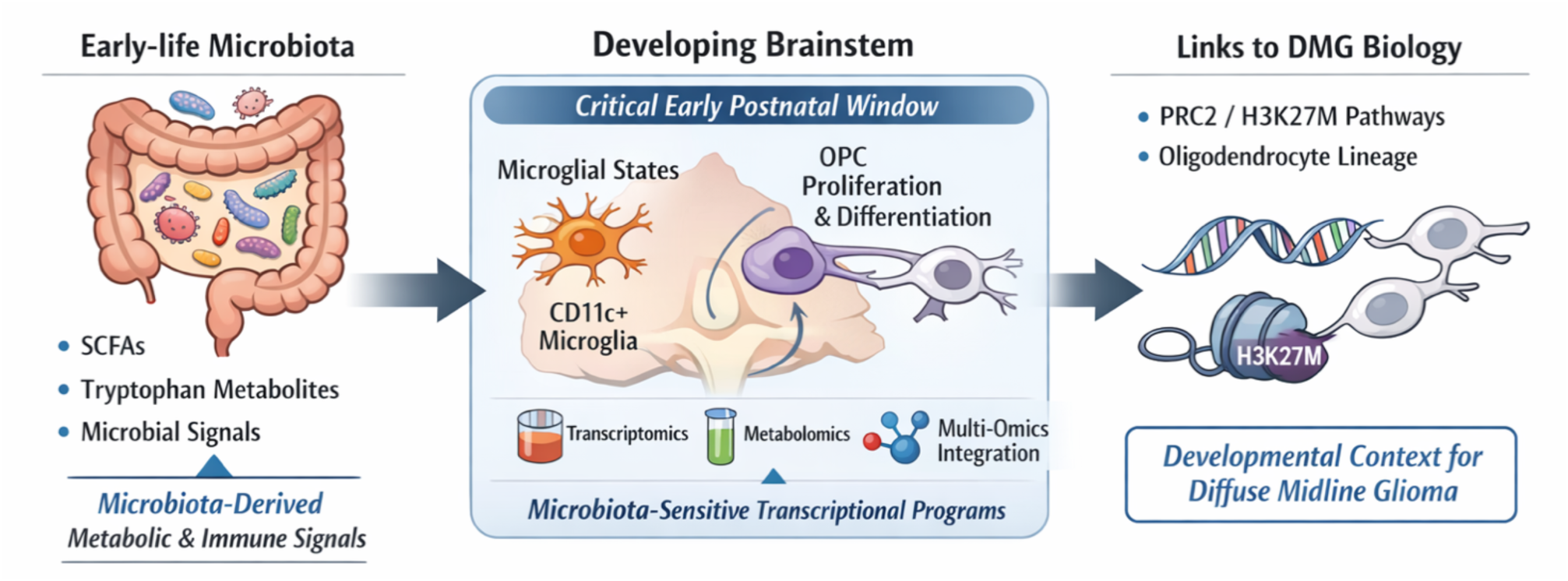
Early-life microbiota provides metabolic and immune cues that shape transcriptional programs in the developing brainstem. Germ-free mice display altered metabolite profiles and transcriptional signatures associated with CD11c⁺ microglia and oligodendrocyte lineage progression. These microbiota-sensitive programs overlap with transcriptional states observed in diffuse midline glioma (DMG).

To further explore the metabolic underpinnings of this microglial program, we performed integrated gene–metabolite analysis and found that Spp1, an important trophic factor of CD11c⁺ microglia, was closely associated with NADH, ribulose-5-phosphate, UDP, and GDP. These metabolites converge on pathways that support cellular energy metabolism, redox balance, and biosynthetic activity, functions essential for sustaining the secretory and supportive roles of Spp1⁺ microglia. The link to NADH suggests that these cells rely on high mitochondrial activity to fuel the production and release of trophic factors (61), while the association with ribulose-5-phosphate implicates the pentose phosphate pathway in maintaining NADPH-dependent antioxidant capacity and nucleotide synthesis (62). UDP and GDP intermediates, central to glycosylation and purine metabolism, may further facilitate protein trafficking and cytokine signalling that influence oligodendrocyte lineage maturation (63, 64). Together, these findings suggest that Spp1⁺ microglia are metabolically primed to support OPC differentiation, coupling energetic and redox homeostasis to the production of pro-myelinating factors such as SPP1 and IGF1 (22, 49, 59, 65). In the brainstem of germ-free animals, perturbation of these metabolite networks may impair the bioenergetic and anabolic capacity of Spp1⁺ microglia, thereby limiting their ability to promote OPC maturation, a mechanism that may be reactivated or co-opted within the tumour microenvironment of DMG. Indeed, DMG tumour cells may exploit this same Spp1⁺ microglial program characterized by elevated oxidative and anabolic metabolism to sustain progenitor proliferation and immune evasion (28, 66). This convergence between developmental and neoplastic microglial states underscores how disruptions in early-life metabolic and microbial cues can create a permissive environment for tumour-promoting neuroimmune interactions (Figure 7).

### Relationship to Disease

The timing of microbiota-associated transcriptional and metabolic differences coincides with periods of rapid pontine growth and myelination in humans(7, 8), raising the possibility that transient microbial perturbations in early life such as antibiotic exposure, caesarean delivery, or dietary factors may influence microbiota-related neurodevelopmental programs. The observed overlap between early postnatal brainstem transcriptional profiles in germ-free animals and gene signatures reported in diffuse midline glioma suggests that early-life environmental perturbations may be associated with persistence of progenitor-related molecular programs. While these findings do not indicate oncogenic transformation, they support the concept that early postnatal developmental stages represent periods of heightened sensitivity during which microbiota-associated signals may exert lasting influences on myelination, neuroimmune maturation, and disease-relevant pathways (Figure 7) (34).

The developmental link between the gut microbiota, CD11c⁺ microglia, and OPC maturation suggests a novel axis for therapy in midline gliomas. Preclinical and clinical evidence in adult gliomas shows that the gut microbiota can influence tumour growth, immune activation/suppression, as well as response to standard therapy (67). Antibiotics can exacerbate tumour progression (68), whereas defined microbial or dietary intervention, such as the ketogenic diet, remodel the microbiota, increase *Akkermansia abundance*, increase butyrate levels, and shift microglia toward a tumour-inhibitory phenotype; notably, this antitumour effect is lost in germ-free or antibiotic-treated conditions and rescued by butyrate or defined consortia (*Akkermansia, Roseburia*)(69–71). A complementary therapeutic avenue lies in targeting one-carbon metabolism: our data show altered SAM biology in the germ-free brainstem during the same developmental stage marked by PRC2 gene dysregulation. In H3K27M DMG, restricting methionine or inhibiting SAM synthesis through MAT2A suppresses tumour growth by restoring PRC2-dependent chromatin repression. Together, these strands argue for microbiota-informed interventions that both re-establish SCFA-driven microglial homeostasis and tune methyl-donor metabolism to normalize PRC2 activity through defined diets (e.g., ketogenic or methionine-restricted), pre-or probiotics (*Akkermansia, Roseburia*), or faecal microbiota transplant strategies embedded within biomarker-driven clinical trials.

### Limitations

While this study identifies microbiota-associated transcriptional and metabolic programs during early postnatal brainstem development, several important limitations should be acknowledged. First, our analyses are based on bulk RNA sequencing and metabolomic profiling of whole brainstem tissue, which precludes definitive assignment of observed molecular changes to specific cell types. Although we infer involvement of microglia and oligodendrocyte lineage cells based on gene set enrichment and integrated pathway analyses, single-cell and spatially resolved approaches will be required to validate cell-type–specific responses to microbial cues.

Second, our study focuses on two early postnatal time points and does not include additional developmental stages or comparisons across multiple brain regions. As such, we cannot exclude the possibility that microbiota-associated transcriptional and metabolic programs extend beyond the brainstem or persist outside the postnatal stages examined. Third, while germ-free mice provide a powerful system for identifying microbiota-sensitive developmental processes, they represent an extreme perturbation that may not fully reflect physiological microbial variation in humans.

In addition, although our integrated analyses highlight metabolic pathways intersecting with those implicated in oligodendrocyte maturation, microglial function, and diffuse midline glioma, we do not establish causal roles for specific microbial taxa or metabolites. Future studies employing gnotobiotic models colonized with defined microbial consortia, targeted metabolite supplementation, or genetic and pharmacological perturbations will be necessary to test sufficiency and mechanism. Finally, our study does not directly model tumorigenesis, and therefore the relevance of microbiota-sensitive developmental programs to diffuse midline glioma remains inferential. Extending these findings to disease-relevant models and human cohorts will be essential to establish translational significance.

## Conclusion

Our findings reveal that the early-life gut microbiota programs the metabolic and immune environment of the developing brainstem through coordinated regulation of microglia and oligodendrocyte lineage cells. The absence of microbial cues locks the brainstem into an immature, proliferative state characterized by PRC2-linked transcriptional dysregulation and metabolic depletion, recapitulating features of diffuse midline glioma. By establishing a microbiota–microglia– oligodendrocyte axis that governs postnatal myelination, this study identifies the gut microbiota as a previously unrecognised environmental determinant of brainstem development. These insights, in addition to providing a resource for investigators interested in brainstem early life development suggest that interventions targeting microbial composition or metabolite production could represent tractable strategies to promote healthy neurodevelopment and potentially mitigate vulnerability to brainstem tumours.

## Methods

### Animals

C57BL/6 breeding pairs (Taconic Biosciences) were mated, with later generations employed in experiments. Germ-free and conventionally raised mice were housed in breeding pairs for two weeks. Afterwards, females were individually housed until birth (postnatal day 0 or P0). Mice were maintained under standard laboratory conditions on a 12-hour light/dark cycle with controlled temperature and humidity (21 ± 1°C and 55%–60%), with *ad libitum* access to standard rodent chow (Special Diet Services, #801010) and autoclaved water. A maximum of 2 mice/litter were used for experimental analyses. All animal procedures were approved by the Animal Experimentation Ethics Committee of University College Cork and the Health Products Regulatory Authority (HPRA AE19130 P047/118), and adhered to the European Directive 2010/63/EC, meeting the requirements of the S.I N° 543 of 2012.

### Sex as a biological variable

All experiments were performed in male mice to minimize biological variability and adhere to the 3Rs principle of animal research. Diffuse midline glioma (DMG) shows no known sex difference in incidence, molecular subtype, or patient outcome(72, 73). Inclusion of both sexes would have required doubling cohort sizes without a clear biological rationale, thereby increasing animal use unnecessarily. Future studies will determine whether sex-specific microbial or neurodevelopmental differences emerge in this context.

### Brainstem tissue collection

Mice designated for fresh tissue collection were sacrificed by decapitation between 10 am and 12 pm. Brainstem tissue was rapidly dissected on ice and immediately snap-frozen on dry ice. Samples were stored at -80°C until further analysis.

### RNA isolation from brainstem

Total RNA was extracted from the brainstem of P2-and P8-old male mice (n = 8/group/age) using the RNeasy® Plus Universal Mini Kit (Qiagen, #73404) following manufacturer’s instructions. Quality and quantity of the isolated RNA were assessed using a Nanodrop ND-1000 spectrophotometer (Thermo Scientific) and using the Qubit™ RNA IQ Assay Kit (Invitrogen, #Q33221). Samples were kept at -80°C until further analysis.

### RNA-Sequencing

mRNA sequencing was conducted by Azenta Life Sciences (Strand-specific RNA-Seq) using strand-specific RNA-Seq on the Illumina NovaSeq 6000 platform (2×150 bp paired-end configuration) and data quality control as well as FASTAQ-file generation was performed by Azenta Life Sciences. Raw sequencing data was analysed using the rnaseq pipeline (https://nf-co.re/rnaseq/3.18.0/docs/output) from the nf-core framework (v24.10.6)(74) using default parameters. Extensive documentation regarding the software used, their default arguments and specific versions can be found in the official GitHub repository of the pipeline (https://github.com/nf-core/rnaseq/tree/3.18.0?tab=readme-ov-file). Briefly, TrimGalore!, FASTQC and SortMeRNA were used for primer removal, sequence quality check and removal of ribosomal RNA, respectively. Kallisto was used to annotate and quantify transcripts using the *Mus musculus* GRCm39.113 reference genome. Resulting h5 files for each sample were imported into R (v4.5.1) with the RStudio GUI (v2023.12.1 Build 402) using the tximport (v1.36.1) package(75).

### Differential gene expression and enrichment analyses

The resulting count data was filtered to retain genes with ≥10 counts in ≥75% of samples, and remaining counts were rounded to integers for DESeq2 analysis.

Differential expression analysis was performed using DESeq2 with the design formula (∼ age + microbiota status + age:microbiota status). Log2 fold changes were shrunk using the apeglm method. Five contrasts were tested: microbiota status at post-natal day (PND) 2 and PND8, age effects in conventional and germ-free mice, and age-microbiota status interactions. P-values were adjusted for multiple testing using the Benjamini-Hochberg (BH) procedure to control the false discovery rate (FDR), with a P-adjusted (*q*) value threshold of 0.05(76) Genes with adjusted *p-value≤0.05* and |log2FC| ≥1 were considered differentially expressed, unless otherwise stated. For targeted enrichment analysis, two complementary approaches on significantly differentially expressed genes (*p<0.05*) were performed from RNA-seq data. First, we compiled manually curated gene sets through literature-based data mining, in PubMed (http://www.ncbi.nlm.nih.gov/pubmed) and manual annotation of candidate genes relevant to specific biological pathways (Supplemental Data 1). Second, relevant gene sets from established Gene OntologyDatabase(77) terms were selected in the Mouse Genome Informatics database (https://www.informatics.jax.org/) to identify candidate genes. Hypergeometric enrichment testing was performed for both approaches using the base R phyper function to identify significantly enriched pathways (*p<0.05*). Principal component analysis (PCA), volcano and heatmap plots were plotted using R. To examine global transcriptomic differences, permutational multivariate analysis of variance (PERMANOVA) was performed using the adonis2 function from the vegan R package (v2.7-1). Euclidean distances were calculated from variance-stabilized count data, and PERMANOVA tested the effects of microbiota status, age, and their interactions using 1,000 permutations.

### Untargeted metabolomic analyses

Brainstems from male mice at P2 and P8 were analysed for untargeted metabolomics (n = 8/group/age). Untargeted metabolomic analyses was conducted by MS-Omics (Copenhagen). Sample preparation involved acidification with hydrochloric acid followed by the addition of deuterated internal standards. Samples underwent randomized analysis using an ultra-performance liquid chromatography system (Vanquish, Thermo Fisher Scientific) connected to a high-resolution quadrupole-orbitrap mass spectrometer (Orbitrap Exploris 240 MS, Thermo Fisher Scientific). The mass spectrometer utilized electrospray ionization with alternating positive and negative ion modes through polarity switching. Quality control samples were included in the analytical sequence. Peak detection and processing were performed using Compound Discoverer software (version 3.3, ThermoFisher Scientific) and Skyline software (version 22.1, MacCoss Lab Software). Peak quantification was based on area under the curve measurements.

For metabolite identification, LC-MS/MS data annotation followed established confidence levels. Level 1 identifications (utilizing accurate mass, MS/MS fragmentation patterns, and retention time matching against in-house reference standards run on identical instrumentation) and Level 2a identifications (employing accurate mass and retention time comparison with in-house standards on the same analytical platform) were retained for subsequent analysis. The final metabolite abundance matrix containing Level 1 and 2a annotations underwent centre log-ratio (CLR) transformation to address the compositional nature of multi-omics datasets using the vegan package in R(78).

Global metabolomic differences were assessed using PERMANOVA with the adonis2 function from the vegan R package. Euclidean distances were calculated from CLR-transformed metabolomics data, testing effects of microbiota status, age, and their interaction using 1,000 permutations. Differential abundance analysis was calculated for each metabolite in the final CLR-transformed data were assessed by fitting the following linear models:metabolite ∼ microbiota status + age or metabolite ∼ microbiota status by age. Untargeted enrichment analysis was performed on differentially abundant metabolites between germ-free and conventional mice at P2 (*p*<0.05) using MetaboAnalyst with the murine KEGG library as a reference(79).

Statistical and bioinformatics analyses were carried out in R statistical software (v4.5.1) using the RStudio interface (v2023.12.1 Build 402). Principal component analysis was applied to the CLR-transformed data following established compositional data analysis methods.

### Integrated multi-omics pathway analysis

For untargeted integrated pathway analysis, significantly altered metabolites and differentially expressed genes in the brainstem of conventional and germ-free mice at P2 were combined for integrated analysis (*p<0.05*). MetaboAnalyst software (v6.0) was employed with standard parameter configurations to interrogate all available KEGG pathway databases(79) for identification of biologically relevant pathways via multi-omics investigation(80) The analytical approach evaluates both the enrichment ratio of pathway components and the topological importance/centrality in the pathway to calculate statistical significance and pathway impact scores(79).

For targeted integrated analysis at P2, gene symbols from the filtered and processed RNA-seq data were mapped to Ensembl IDs using the org.Mm.eg.db package (v3.21.0), then converted to KEGG ortholog (KO) identifiers via EntrezID and EC number mappings using KEGG REST API (https://rest.kegg.jp/). RNA-seq count data for mapped genes were aggregated by KO and centre log-ratio (CLR) transformed. Metabolites were mapped to KEGG compound identifiers using MetaboAnalyst reference data, and CLR-transformed metabolomics data were filtered to retain only metabolites with valid KEGG IDs. For each KO-metabolite pair linked within a KEGG pathway (based on KEGG pathway annotations), Pearson correlations were calculated separately within conventional and germ-free groups. Differences in correlation coefficients between microbiota states were assessed using Fisher’s z-transformation, with p-values adjusted for multiple testing using the Benjamini-Hochberg method. Associations with adjusted *p<0.05* were considered statistically significant. For targeted integrated analysis for CD11c^+^ microglia signature at P2, direct gene-metabolite correlation analysis was performed instead of KEGG pathway-based integration. This alternative approach was necessary since insufficient KEGG ortholog annotations were identified for CD11c+ signature genes. CLR-transformed gene expression and CLR-transformed metabolite abundances were directly correlated using Pearson correlations calculated separately for conventional and germ-free groups. Differential associations between microbiota status were assessed using linear models with interaction terms (metabolite ∼ gene × microbiota status), comparing full models against reduced models (metabolite ∼ gene + microbiota status) via ANOVA. P-values were adjusted for multiple testing using the Benjamini-Hochberg method. Associations with overall correlation adjusted *q<0.1* and interaction adjusted *q<0.05* were considered significant.

### Single-cell RNA-sequencing data analysis

A 4,058 × 23,686 gene-level transcripts-per-million (TPM) matrix for the K27M cohort and its metadata were obtained from the Gene Expression Omnibus (accession **GSE102130**) and imported into an R-environment (version 4.5.1). Cells were filtered by their previously defined annotation [https://doi.org/10.1126/science.aao4750], and immune cells from two patients (BCH836 and BCH1126) with more than three immune cells from the pons were retained for downstream analysis (92 cells total). The raw TPM values were stored in a

SingleCellExperiment object (SingleCellExperiment, v1.30.1) [https://doi.org/10.1038/s41592-019-0654-x] and natural-log-transformed (logcounts = log1p(TPM)). Genes with mitochondrial (“MT-”) and ribosomal (“RPL”, “RPS”) prefixes were removed. All downstream analyses used the logcounts assay unless explicitly stated.

Gene-level variance modelling was performed with modelGeneVar (scran, v1.36.0) [https://doi.org/10.12688/f1000research.9501.2] while blocking for patient, and the top 2,000 highly variable genes (HVGs) were selected using getTopHVGs from the same package. Principal component analysis (PCA) was computed followed by batch correction across patients using mutual nearest neighbors with fastMNN (batchelor, v1.24.0) [https://doi.org/10.1038/nbt.4091] on the HVGs (parameters: *d = 9*, *k = 20*, cosine normalization = TRUE). UMAP embeddings were generated on MNN-corrected principal components with runUMAP (scater, v1.36.0) [https://doi.org/10.1038/s41467-022-29212-9] and visualized with continuous UCell signature scores calculated by ScoreSignatures_UCell (UCell, v2.12.0) [https://doi.org/10.1016/j.csbj.2021.06.043]. Cells were scored against four a priori microglial signatures using default UCell parameters (**KS+**: B4GALT1, B3GALT4, B3GNT2, CHST1, CHST2, CHST3, CHST4, CHST5, GAL, OGT; **Hoxb8+**: HOXB8, ITGA4, PLBD1, CD300LD, SAMD9L, CCR2, SERPINC1, CYTIP, GPR65, SH3PXD2B; **Arginase+**: ARG1, COLEC12, TRIM6, ORC1, ADAM8, HGF, RHBDL1, CD274, F2R, SAP25; **CD11c/IGF1+**: ITGAX, IGF1, IGF1R, LPL, SPP1, GPNMB, CSF1, FABP5, LGALS1, CLEC7A). For each cell, UCell scores, based on Mann-Whitney U statistics, were computed per signature, and a signature label was assigned as the maximum value among the four scores (ties.method = "first"). Expression values were averaged per label (mean across cells within each group) to obtain a genes × groups matrix. Two versions were prepared: (i) unscaled averages, representing log1p(TPM) means by group (to display absolute magnitude), and (ii) scaled (z-score) values by row-wise normalization across groups. Within each signature, genes were ordered by hierarchical clustering of group means (hclust - stats, v4.5.1), and heatmaps were generated with ComplexHeatmap, v2.24.1[https://doi.org/10.1093/bioinformatics/btw313].

### Statistical analysis

All data, including transcriptomic and metabolomic were analysed using R (v4.5.1) with the RStudio GUI (v2023.12.1 Build 402) as detailed above.

### Data availability

Transcriptomic data have been deposited and will be publicly available at the European Nucleotide Archive with accession number, PRJEB97701, upon publication. Metabolomic dataset have been deposited and will be available on MetaboLights with accession number REQ20250922213321. All statistical analyses are available in Supplemental Data 1.

### Code availability

Original code will be made available upon request.

## Declarations

### Ethics approval

All animal procedures were approved by the Animal Experimentation Ethics Committee of University College Cork and the Health Products Regulatory Authority (HPRA AE19130 P047/118), and adhered to the European Directive 2010/63/EC, meeting the requirements of the S.I N° 543 of 2012.

### Consent for publication

All authors have consented to publication of the submitted manuscript

## Acknowledgements

We extend our appreciation to Patrick Fitzgerald, Colette Manley, Jessica O’Reilly, Laicee Kenny, Frances O’Brien and Dr. Kenneth O’Riordan for their invaluable technical expertise and support.

## Authors information

LK and JFC conceived the study. LK and CCM designed experiments, interpreted the data and wrote the first draft of the manuscript. CCM performed all data analysis with help from LW for metabolomics, BV, JAL and AB for bulk RNA sequencing, and CMKL for integration analysis. CMKL also collected brainstem tissues. JE analysed DMG single cell data. SG, GC. GMM and JFC assisted with data interpretation and critically reviewed the manuscript.

